# Development of *Kaptive* databases for *Vibrio parahaemolyticus* O- and K-antigen serotyping

**DOI:** 10.1101/2021.07.06.451262

**Authors:** Linda van der Graaf – van Bloois, Hongyou Chen, Jaap A. Wagenaar, Aldert L. Zomer

**Affiliations:** Department of Biomolecular Health Sciences, Faculty of Veterinary Medicine, Utrecht University, the Netherlands; WHO Collaborating Centre for Reference and Research on Campylobacter and Antimicrobial Resistance from a One Health Perspective / OIE Reference Laboratory for Campylobacteriosis, Utrecht, the Netherlands; Shanghai Municipal Center for Disease Control and Prevention, No. 1380, Zhong Shan Xi Rd., Shanghai 200336, PR China; Wageningen Bioveterinary Research, Lelystad, the Netherlands

**Keywords:** Vibrio parahaemolyticus, serotyping, whole genome sequencing, O-locus, K-locus

## Abstract

*Vibrio parahaemolyticus* is an important food-borne human pathogen and is divided in 16 O- serotypes and 71 K-serotypes. Agglutination tests are still the gold standard for serotyping, but many *V. parahaemolyticus* isolates are not typable by agglutination. An alternative for agglutination tests is serotyping using whole genome sequencing data. In this study, *V. parahaemolyticus* isolates are serotyped and sequenced, and all known and several novel O- and K-loci are identified. We developed *Kaptive* databases for all O- and K-loci after manual curation of the loci. These *Kaptive* databases with the identified *V. parahaemolyticus* O- and K -loci can be used to identify the O- and K-serotypes of *V. parahaemolyticus* isolates from genome sequences.

## Introduction

*Vibrio parahaemolyticus* (*V. parahaemolyticus*) is an important food-borne human pathogen that naturally inhabits marine environments worldwide, and can cause acute gastroenteritis and septicemia in human (1,2). *V. parahaemolyticus* is typically serotyped on the basis of its heat-stable somatic antigen (O) and its capsular antigen (K) and classified into 16 O-serotypes and 71 K-serotypes.

Serotyping of *V. parahaemolyticus* is important for pathogen detection and epidemiological surveillance. Many *V. parahaemolyticus* serotypes have been identified as pandemic clones, and certain serotypes, for example O3:K6, O1:KUT and O4:K68, are generally considered to be more virulent than others (3,4). Agglutination tests are the gold standard for *V. parahaemolyticus* serotyping but frequently, *V. parahaemolyticus* strains are non-typable with agglutination tests in routine screening (3,5–7). For several pathogenic bacterial species, e.g. several pathogenic *E. coli* serogroups (8) and *Vibrio cholerae* serogroup O1 and O139 (9,10), PCR methods are developed to detect the specific serogroups, but no PCR methods are available for *V. parahaemolyticus* serotyping.

The software tool *Kaptive* was developed for rapid O- and K-loci typing of *Klebsiella* strains from whole genome sequences (11). This tool includes an option to use other, self-created databases. The genomic regions associated with the somatic synthesis locus (O-locus) and capsule synthesis locus (K-locus) for *V. parahaemolyticus* are generally found on the same genomic location (2,12,13), which makes it possible to locate and extract the O- and K-loci nucleotide sequences from whole genome sequencing data of *V. parahaemolyticus* strains. Recently, the algorithm VPsero was developed, which identified serogroup-specific genes as markers *in silico*, but only marker genes for 12 O- and 43 K-serotypes were covered (14).

The aim of this study was to identify all 16 O- and 71 K-loci of *V. parahaemolyticus* by serotyping, sequencing and analyzing in-house isolates and investigating public data of serotyped strains and manually curate reference databases for O- and K-loci, which can be used to determine the O- and K-serotype of *V. parahaemolyticus* strains from whole genome sequencing data.

## Materials and Method

### *Datasets of* V. parahaemolyticus

In this study, 56 *V. parahaemolyticus* isolates were sequenced and serotyped. For whole genome sequencing (WGS), DNA was extracted with TIANamp Bacteria DNA Kit (Tiagen Biotech (Beijing) CO., LTD, Beijing, China) according to the manufacturer’s recommendation and DNA quality control was performed using agarose gel electrophoresis and the Qubit dsDNA HS Assay Kit. DNA libraries and DNA nanoballs (DNB) were constructed on BGISP-100 platform (WuHan MGI Tech Co.,Ltd, Wuhan, China) with input of 150ng DNA. 50 bp single-end reads were generated with BGISEQ-50 sequencer (WuHan MGI Tech Co.,Ltd, Wuhan, China) and assembled with SPAdes v3.12 (15). All sequence reads of strains sequenced in this study are available under accession PRJEB39490 and PRJNA483379 of the ENA short read archive.

Two datasets of public available *V. parahaemolyticus* genomes were included, consisting of 776 genomes from NCBI Genbank (Accession date: 14 December 2017) and 1498 genomes from PATRIC database (16) (Accession date: 23 July 2020) were downloaded.

Taxonomic classification of all genomes was performed by *in silico* 16S DNA analysis with SPINGO v1.3 (17) and genomes with ambiguous 16S results were classified as *Vibrio parahaemolyticus* using KmerFinder v3.2 (18).

### Serotyping of V. parahaemolyticus strains

Serotyping of the 56 in this study sequenced strains was performed by slide agglutination according to the manufacturer’s instructions, using the 65 type-K serum set and 11 type-O serum set (Denka Seiken CO., LTD, Tokyo, Japan).

### Development Kaptive databases

#### I. Identification of O-loci

*V. parahaemolyticus* is divided into 16 O-serotypes (1,2). The O-locus has been defined located between *dgkA* and *gmhD* genes, which encode a diacylglylcerol-kinase and an epimerase respectively (2). With BlastN (minimum coverage of 80% and minimum identity of 80%), the flanking genes of the O-region, *dgkA* and *gmhD*, were searched in all genome sequences. The O-loci sequences were extracted from these sequences manually.

#### II. Identification of K-loci

*V. parahaemolyticus* is divided in 71 K-serotypes (2). The K-locus has been defined located between gene *gmhD* which encodes an epimerase at one side, and at the other side gene *rjg* encoding a metallo-hydrolase (13). With BlastN (minimum coverage of 80% and minimum identity of 80%), the flanking genes of the K-region, *gmhD* and *rjg*, were located in all 1601 genome sequences. In some cases where the *rjg* gene was not found, gene *ugd* or gene *gtaB* was selected as flanking gene. The K-loci sequences were extracted from these sequences manually.

#### III. Development of the *Kaptive* databases

The extracted O- and K-loci sequences were annotated using Prokka v1.13 (19). For annotating the loci, a custom database was used which was built with the annotations of the previous described loci of *V. parahaemolyticus* (1,2). Genes of the O- and K-loci respectively were clustered using Roary with an 80% cutoff on amino acid identity (20). The Roary gene presence absence table was used to create gene presence-absence clusters of the O- and K-loci respectively and the unique gene presence-absence loci were linked to the known serotypes of the genomes.

For the identified O- and K-loci, the nomenclature of the *Klebsiella* capsule synthesis loci (11) was used; each distinct O- and K-locus was designated as OL (O-locus) and KL (K-locus), followed by an unique number. The O- and K-loci of known serotypes were assigned with the same number as the corresponding O- and K-serotypes, e.g. O-serotype 1 is encoded by the OL1 locus etcetera. O- and K-loci with unknown serotypes were assigned with numbers starting from 101 (e.g. OL101 and KL101). Variants of loci with maximal two genes different by insertion or deletion were given the suffix -1. Eight gene patterns of K-Untypable (KUT) strains were found and these patterns are assigned in the k-loci database as KLUT (K-locus UnTypable) followed by subsequent numbers 1-8 Of each loci, one genome was selected as reference and the nucleotide sequences of the O- and K-loci of the selected genomes were added to the *Kaptive* database file and curated manually.

The *V. parahaemolyticus* O-locus and K-locus *Kaptive* databases are available on the following Github page: https://github.com/aldertzomer/vibrio_parahaemolyticus_genomoserotyping and will be added to Kaptive web: https://kaptive-web.erc.monash.edu/

### Comparison of genes in O- and K-loci

The reference genomes of the identified O- and K-loci were aligned using Parsnp v1.2 (21), recombination regions were filtered using Gubbins v2.3.4 (22) and the tree was built with FastTree v2.1.8 (23). The tree was visualized with iTol v6.5.4 (24) and Multi Locus Sequence Typing (MLST) was performed in silico with the MLST software (Seemann) using PubMLST typing schemes (26). Gene cluster comparisons of the O- and K-loci are made with Clinker v0.0.37 (27).

### Performance of Kaptive databases compared to VPSero

Of the total set of 831 selected genomes from Genbank (n=775) and in-house sequenced genomes (n=56), 436 genomes had a known O-serotype and 294 genomes had a known K-serotype, whereas the PATRIC genomes had all unknown O- and K-loci serotypes. The O- and K-loci of the genomes with the known serotypes were all serotyped by the newly developed *Kaptive* databases and VPSero (14). Furthermore, performance of the *Kaptive* databases was tested and compared with VPSero by using the VPSero sequence data collection, deposited into CNGB Sequence Archive project CNP0000343 described by Bian *et al* (14).

## Results and Discussion

### Development of Kaptive databases for V. parahaemolyticus O- and K-loci

The study started with a set of 2330 *V. parahaemolyticus* genomes, consisting of 56 in this study sequenced genomes, 776 genomes downloaded from NCBI GenBank and 1498 from the PATRIC database. The genomes that were *in silico* identified as *V. parahaemolyticus* with 16S and KmerFinder analysis were included. For the genomes downloaded from NCBI GenBank and PATRIC database, only genomes that contained the complete loci on one contig were included. This selection ended-up with a total set of 1601 genomes, consisting of 56 in-house sequenced strains, 775 from Genbank and 770 genomes from PATRIC database. For the development of the O- and K-loci databases, only genomes that contained both flanking genes of the loci on one contig were used. No isolate or genome were available for K-serotypes 26 and 71, and these serotypes were therefore not included in the database.

The selected genomes that were used as reference for each O- and K-loci are shown in Supplemental Table 1. For each O-serotype, unique loci were identified, except for serotype O3 and O13 genomes, since the loci of these serotypes contained the same genes (1). Therefore, loci of serotype O3 and O13 could not be distinguished based on gene presence and are assigned in the *Kaptive* O-database as “O3_or_O13”. It is possible that a second cluster modifies the O-antigen, similar to what has been described for *Shigella* (28), however an O-modifying locus has not been described in literature for O13.

Eight gene patterns of K-UnTypable (KUT) strains previously described were found and these patterns are assigned in the *Kaptive* database as K-Locus UnTypable (KLUT) followed by subsequent numbers 1-8 (Supplemental Table 1).

For both O- and K-loci, not all gene patterns could be assigned to specific serotypes, because these gene patterns were found in genomes with unknown serotypes from NCBI and PATRIC databases. These unknown gene patterns were assigned for O-loci as OL101 and for K-loci as KL101-KL157 (Supplemental Table 1).

For some O- and K-serotypes, variants in the loci gene patterns by insertion of deletion of maximal two genes were found, and therefore, these serotypes have two gene patterns included in the database, distinguished with the addition of “-1” to the locus name (OL1, OL4, OL7 and KL20, KL30, KL68, KLUT4) (Supplemental Figures 1 and 2).

Examination of the phylogenetic tree of the reference isolates showed the O- and K-loci are not associated with MLST sequence types, e.g. six different K-loci and four different O-loci are found in ST3 reference isolates (Figure 1).

**Figure 1.**
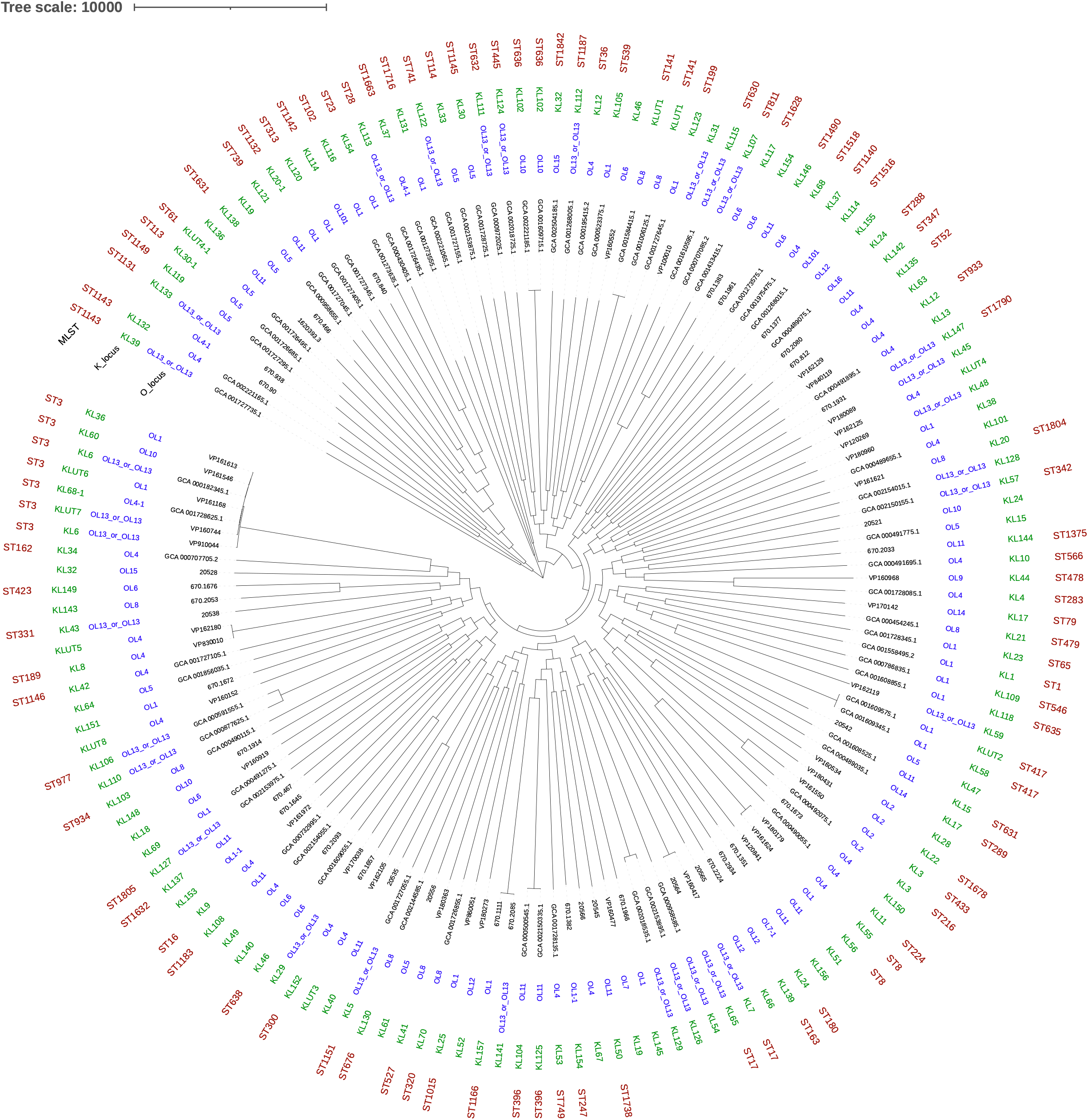
Phylogenetic tree based on core genome analysis of O- and K-representative isolates, including the O- and K-loci, and MLST STs.

### Performance of Kaptive databases

#### Genome collection of this study

To investigate the performance of the newly developed *Kaptive* databases, 436 genomes with a known O-serotype and 295 genomes with a known K-serotype were used from our genome collection. Of the 436 known O-serotypes, 398 O-serotypes were correctly identified (Figure 2A, Supplemental Table 1) and 265 of the 295 known K-serotypes were correctly identified (Figure 2B, Supplemental Table 1).

**Figure 2.**
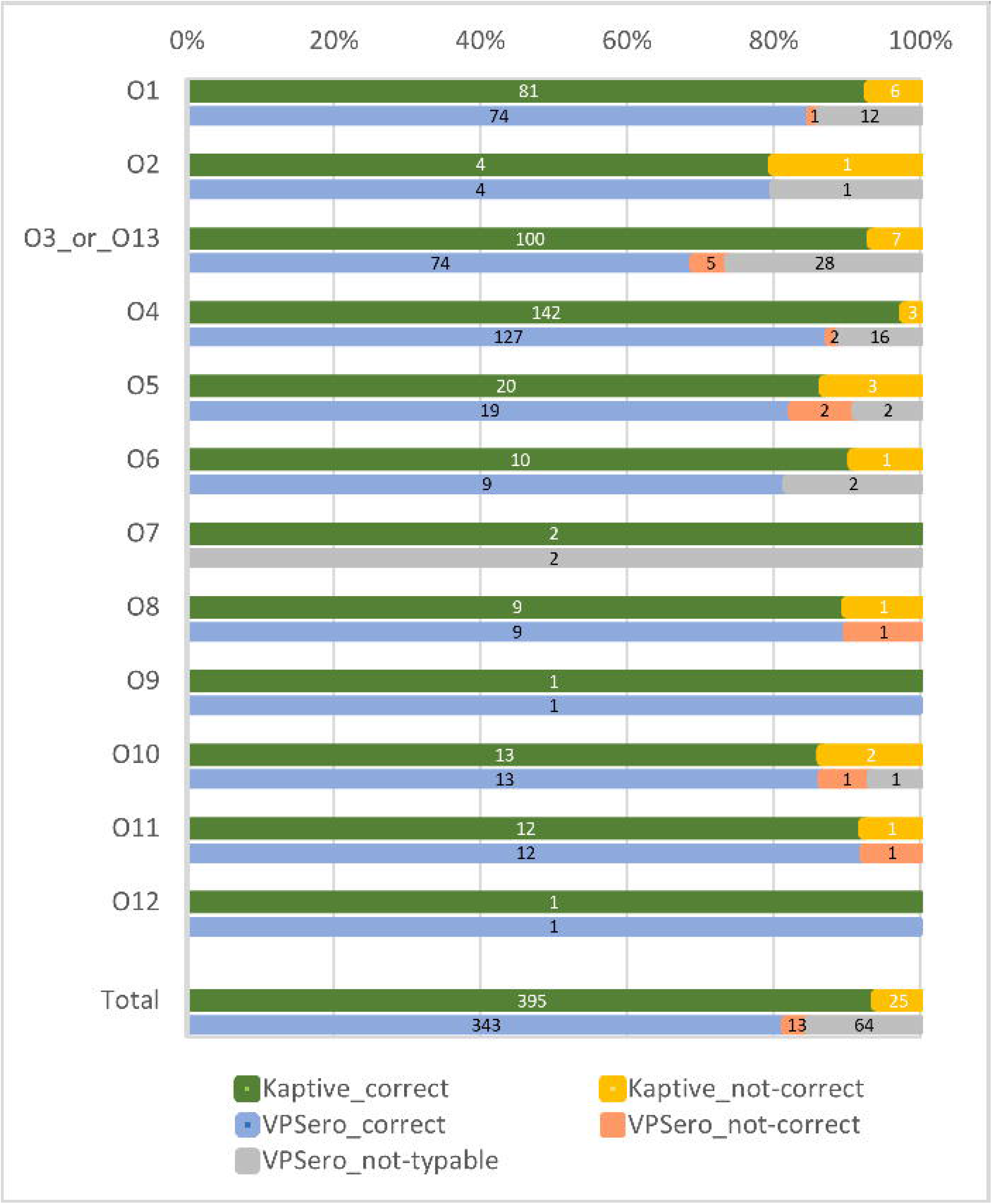
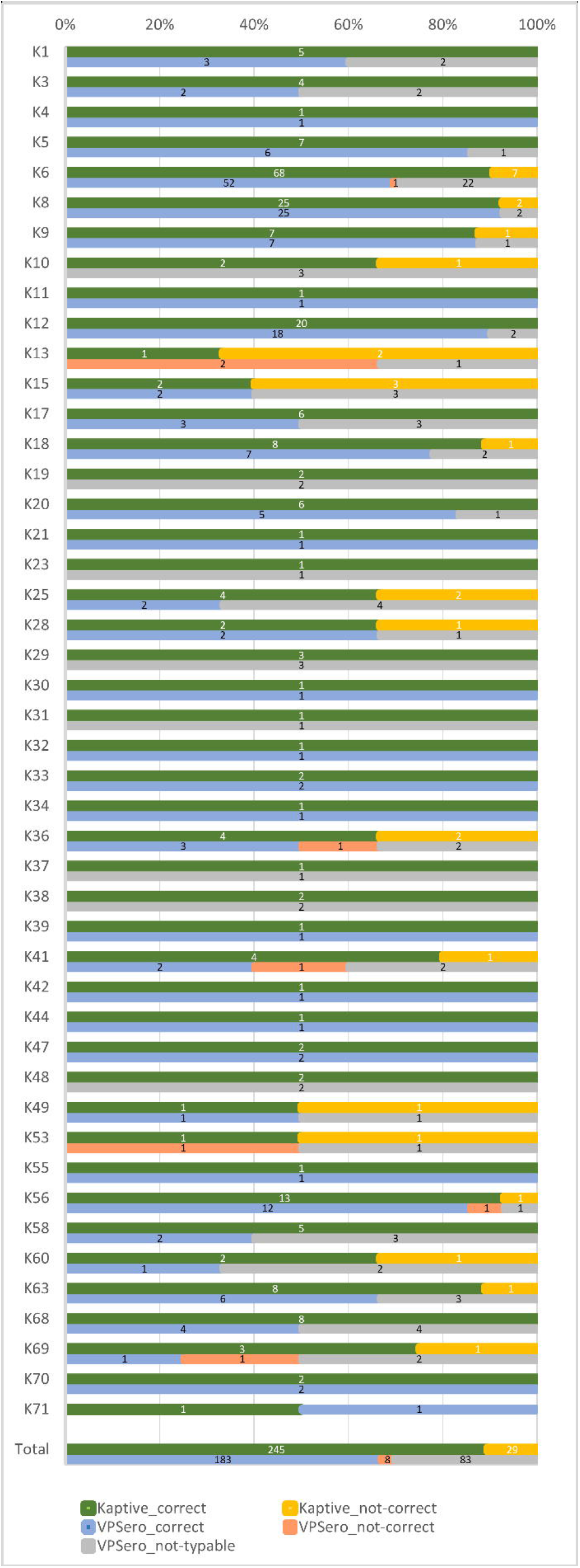
Performance of the *Kaptive* database and VPSero on genomes with known (A) O- and (B) K-serotypes. In green the percentages of correctly typed genomes with *Kaptive* databases, in orange the percentages of incorrectly typed genomes with *Kaptive* databases, in blue the percentages of correctly typed genomes with VPSero, in red the percentages of incorrectly typed genomes with VPSero, in grey the percentages of the with VPSero not-typable genomes. The numbers in the bars represent the number of genomes.

The tool VPSero was developed for 43 K- and 12 O-serotypes (14), and covers part of the known 16 O-serotypes and 71 K-serotypes. The performance of VPSero was tested with a set of 420 genomes for O-serotyping and 274 genomes for K-serotyping from our genome collection, consisting of genomes with known serotypes of the 12 and 46 O- and K-serotypes respectively which are included in VPSero. Of this set, 343 out of 420 genomes with known O-serotypes were correctly identified (Supplemental Figure 2A, Supplemental Table 1) and 183 out of 274 genomes with known K-serotypes (Supplemental Figure 2B, Supplemental Table 1). The newly developed *Kaptive* databases outcompete VPSero with 395/420 correctly identified O-serotype genomes and 254/274 correctly identified K-serotype genomes (Supplemental Figure 2AB, Supplemental Table 1).

#### Sequence collection of VPsero

In the publication of VPsero, intact LPS and CPS gene clusters (LPS/CPSgc) were identified and extracted, and deposited in CNGB Sequence Archive project CNP0000343. These gene clusters did not contain the full O-serotype loci of the newly developed O-serotype *Kaptive* database, and therefore, these sequence data collection could not be used to test the performance of the O-serotype *Kaptive* database, the K-serotype loci were available however. The LPS/CPSgcs were tested with both *Kaptive* K-database and VPSero, and results are listed in Supplemental Table 2. One K58 gene cluster was identified as K6 by both tested tools, possibly containing a wrong serotype in the CNGB database. Of the remaining 56 gene clusters with known K-serotypes, seven were not typable by VPSero and two were mis-identified by *Kaptive* databases.

The mis-identified O- and K-serotypes from our collection were all genomes downloaded from NCBI. Since we only have the genomes and not the strains, we cannot check if these strains were serotyped correctly. O-serotype O14 has been recently described (2) and only one genome with this O-serotype is available. In our dataset of 831 genomes, 14 NCBI genomes with serotype O5 are identified with the *Kaptive* database as the new serotype O14. It is very likely that these genomes are mis-serotyped, because the serum for serotype O14 was not available when these genomes were serotyped.

For several K-serotypes, only one genome sequence was available and we could not determine if there is variation in these K-loci gene content. Furthermore, several of the genomes sequenced in this study were sequenced with a 50-bp single-end BGI sequencer, resulting in a higher number of short contigs, therefore for several of these genomes, the K-locus was not assembled on one single contig. For these genomes, the contigs with flanking genes were selected and concatenated manually. It is possible that some genes are missing in these K-locus reference sequences, although these likely represent repeat sequences. If more *V. parahaemolyticus* genome sequences with closed K-loci become available, the *Kaptive* K-serotype database will be updated with the closed K-locus sequence of these K-serotypes.

## Conclusion

The in this study developed *Kaptive* databases with the identified 16 O- and 71 K -loci can be used to identify the O- and K-serotypes of *V. parahaemolyticus* isolates from whole genome sequencing data. The variation of K-antigen loci is much higher than expected as we identified 57 new K-locus variants.

## Author Statements

### Author contributions

Designed the study; L.G., J.W., A.Z. Performed the experiments; L.G., H.C., A.Z. Analyzed the data; L.G., A.Z. Writing the paper; all authors.

### Conflicts of interest

The authors declare that they have no competing interests.

### Funding information

Sequencing of isolates was supported by the National Science and Technology Major Project of China (No. 2018ZX10305409-003).

## Acknowledgements

We thank the Kaptive team (Thomas Stanton, Kelly Wyres, Ryan Wick and Kathryn Holt) for the undertaking to include the *Vibio parahaemolyticus* databases to Kaptive web.

## Abbreviations

KL: K-locus
KUT: K-Untypable
KLUT: K-locus Untypable
LPS/CPSgc: LPS and CPS gene clusters
MLST: Multi Locus Sequence Typing
OL: O-locus
*V. parahaemolyticus*: Vibrio parahaemolyticus
WGS: whole genome sequencing

## Figures and Tables

**Supplemental Figure 1**. Gene cluster comparisons of the identified O-loci. All similar gene homologs have been assigned an unique color and are linked with similar color boxes. Hypothetical proteins are assigned with an “1”.

**Supplemental Figure 2**. Gene cluster comparisons of the identified K-loci. All similar gene homologs have been assigned an unique color and are linked with similar color boxes. Hypothetical proteins are assigned with an “1”.

**Supplemental Table 1**. Selected reference genomes used for the O- and K-loci *Kaptive* databases. Included are the numbers of genomes with known O- and K-serotypes and numbers of correctly identified genomes with *Kaptive* databases and VPSero.

**Supplemental Table 2**. Performance of *Kaptive* databases and VPSero on sequence collection of VPSero.

## Notes

**Data statement:** All supporting data, code and protocols have been provided within the article or through supplementary data files. The supplementary tables and supplementary figures are available with the online version of this article.

### Competing Interest Statement

The authors have declared no competing interest.

### Summary of Updates

We included the V. parahaemolyticus genomes of Patric database to the analysis. We changed the nomenclature of our databases to the same as the Klebsiella capsule synthesis loci. We included an analysis of the performance of the new Kaptive databases compared to the recently developed algorithm VPSero.

