## Supplementary figures and images for "Development of *Kaptive* databases for *Vibrio parahaemolyticus* O- and K-antigen serotyping"

### Supplemental Figure 1

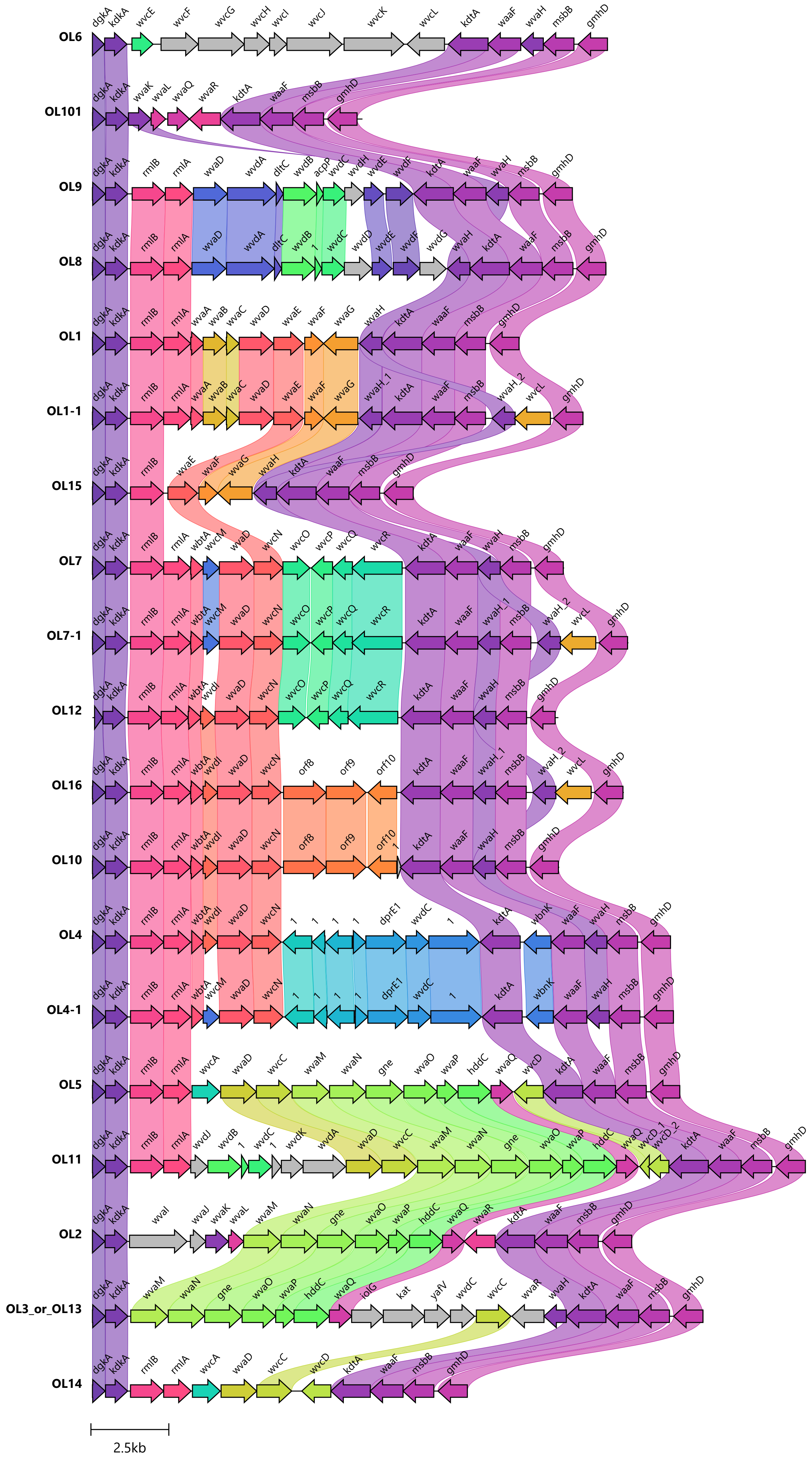

### Supplemental Figure 2

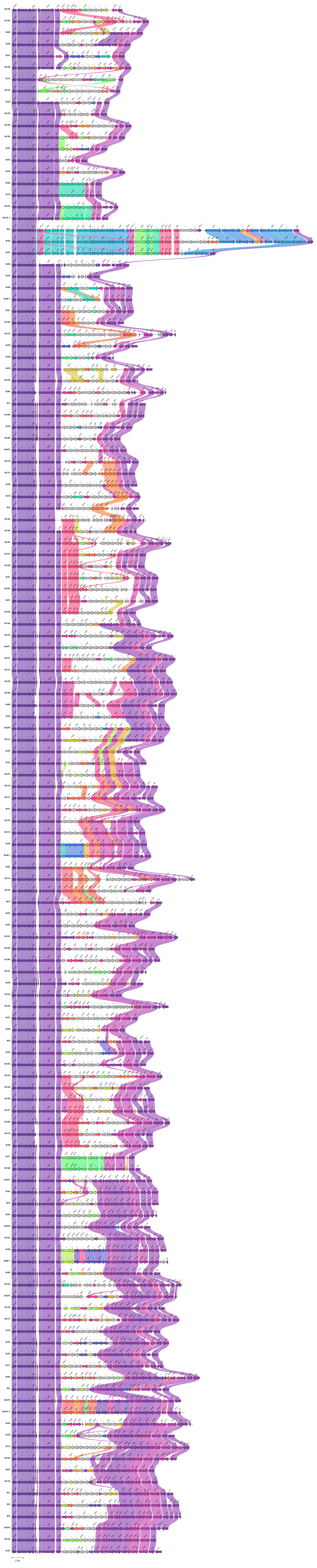
