## Supplemental Table 1 for "Development of *Kaptive* databases for *Vibrio parahaemolyticus* O- and K-antigen serotyping"

| O-serotype | O-locus name | | Reference genom | Sequences obtained from | Accession number | Total genomes with this O-serotype | Kaptive_Correct | VPSero_Correct |
|---|---|---|---|---|---|---|---|---|
| 1 | OL1 | in database | GCA_000430405 | NCBI | | 89 | 82 | |
| 1 | OL1-1 | in database | 670.1382 | PATRIC | | | | |
| 2 | OL2 | in database | GCA_000492075 | NCBI | | 8 | 7 | |
| 3 or 13 | OL3_or_OL13 | in database | GCA_000182345 | NCBI | | 109 | 98 | |
| 4 | OL4 | in database | GCA_000195415 | NCBI | | 145 | 141 | |
| 4 | OL4-1 | in database | GCA_001273555 | NCBI | | | | |
| 5 | OL5 | in database | GCA_000491775 | NCBI | | 39 | 22 | |
| 6 | OL6 | in database | GCA_001609055 | NCBI | | 12 | 11 | |
| 7 | OL7 | in database | VP160477 | this study | GCA_905331675 | 1 | 1 | |
| 7 | OL7-1 | in database | 670.2934 | PATRIC | | | | |
| 8 | OL8 | in database | GCA_001006125 | NCBI | | 10 | 9 | |
| 9 | OL9 | in database | VP160968 | this study | GCA_905331755 | 1 | 1 | |
| 10 | OL10 | in database | GCA_001609715 | NCBI | | 16 | 14 | |
| 11 | OL11 | in database | GCA_001608525 | NCBI | | 15 | 13 | |
| 12 | OL12 | in database | VP180273 | this study | | 1 | 1 | |
| 14 | OL14 | in database | GCA_000489035 | NCBI | | 1 | 1 | |
| 15 | OL15 | in database | GCA_002504185 | NCBI | | 1 | 1 | |
| 16 | OL16 | in database | GCA_000489075 | NCBI | | 1 | 1 | |
| | | | | | | 449 | 403 | |
| **Serotype Unknown/New** | | | | | | | | |
| | OL101 | in database | 670.840 | PATRIC | | | | |

| K-serotyp | K-locus name | | Reference genome | Sequences obtained from | Accession number | k-serotype | Kaptive_Correct | VPSero_Correct | | |
|---|---|---|---|---|---|---|---|---|---|---|
| 1 | KL1 | in database | GCA_001558495 | NCBI | | 5 | 5 | 3 | n.a. | not applicable |
| 2 | | does not exist | | | | | | | | |
| 3 | KL3 | in database | VP161550 | this study | GCA_905331795 | 4 | 4 | 2 | | |
| 4 | KL4 | in database | GCA_001728085 | NCBI | | 1 | 1 | 1 | | |
| 5 | KL5 | in database | GCA_001727055 | NCBI | | 7 | 7 | 6 | | |
| 6 | KL6 | in database | VP910044 | this study | GCA_905332005 | 75 | 68 | 52 | | |
| 7 | KL7 | in database | VP160417 | this study | GCA_905331685 | 1 | 1 | n.a. | | |
| 8 | KL8 | in database | VP830010 | this study | GCA_905331985 | 27 | 25 | 25 | | |
| 9 | KL9 | in database | VP161972 | this study | GCA_905331865 | 8 | 7 | 7 | | |
| 10 | KL10 | in database | GCA_000491695 | NCBI | | 3 | 2 | 0 | | |
| 11 | KL11 | in database | GCA_000490055 | NCBI | | 1 | 1 | 1 | | |
| 12 | KL12 | in database | VP840119 | this study | GCA_905331995 | 20 | 20 | 18 | | |
| 13 | KL13 | in database | GCA_000491895 | NCBI | | 3 | 1 | 0 | | |
| 14 | | does not exist | | | | | | | | |
| 15 | KL15 | in database | GCA_000491775 | NCBI | | 5 | 2 | 2 | | |
| 16 | | does not exist | | | | | | | | |
| 17 | KL17 | in database | VP170142 | this study | GCA_905331935 | 6 | 6 | 3 | | |
| 18 | KL18 | in database | VP160919 | this study | GCA_905331745 | 9 | 8 | 7 | | |
| 19 | KL19 | in database | GCA_000958655.1 | NCBI | | 2 | 2 | 0 | | |
| 20 | KL20 | in database | VP161621 | this study | GCA_905331805 | 6 | 6 | 5 | | |
| 20 | KL20-1 | in database | GCA_001727405 | NCBI | | | | | | |
| 21 | KL21 | in database | GCA_000454245 | NCBI | | 1 | 1 | 1 | | |
| 22 | KL22 | in database | VP180431 | this study | GCA_905331925 | 1 | 1 | n.a. | | |
| 23 | KL23 | in database | GCA_001728345 | NCBI | | 1 | 1 | 0 | | |
| 24 | KL24 | in database | 20521 | this study | GCA_905331545 | 1 | 1 | n.a. | | |
| 25 | KL25 | in database | VP860051 | this study | GCA_905331975 | 6 | 4 | 2 | | |
| 26 | | no isolate/genome with this K serotype | | | | | | | | |
| 27 | | does not exist | | | | | | | | |
| 28 | KL28 | in database | VP160534 | this study | GCA_905331725 | 3 | 2 | 2 | | |
| 29 | KL29 | in database | VP170038 | this study | GCA_905331875 | 3 | 3 | 0 | | |
| 30 | KL30 | in database | GCA_001728725 | NCBI | | 1 | 1 | 1 | | |
| 30 | KL30-1 | in database | GCA_001726685 | NCBI | | | | | | |
| 31 | KL31 | in database | VP100010 | this study | GCA_905331655 | 1 | 1 | 0 | | |
| 32 | KL32 | in database | 20528 | this study | GCA_905331555 | 1 | 1 | 1 | | |
| 33 | KL33 | in database | GCA_002153875 | NCBI | | 2 | 2 | 2 | | |
| 34 | KL34 | in database | GCA_000707705 | NCBI | | 1 | 1 | 1 | | |
| 35 | | does not exist | | | | | | | | |
| 36 | KL36 | in database | VP161613 | this study | GCA_905331785 | 6 | 4 | 3 | | |
| 37 | KL37 | in database | GCA_001975475 | NCBI | | 1 | 1 | 0 | | |
| 38 | KL38 | in database | VP180960 | this study | GCA_905331965 | 2 | 2 | 0 | | |
| 39 | KL39 | in database | GCA_001727735 | NCBI | | 1 | 1 | 1 | | |
| 40 | KL40 | in database | 20535 | this study | GCA_905331575 | 1 | 1 | n.a. | | |
| 41 | KL41 | in database | VP180363 | this study | GCA_905331945 | 5 | 4 | 2 | | |
| 42 | KL42 | in database | GCA_001727105 | NCBI | | 1 | 1 | 1 | | |
| 43 | KL43 | in database | 20538 | this study | GCA_905331585 | 1 | 1 | n.a. | | |
| 44 | KL44 | in database | VP160968 | this study | GCA_905331755 | 1 | 1 | 1 | | |
| 45 | KL45 | in database | VP180089 | this study | GCA_905331955 | 1 | 1 | n.a. | | |
| 46 | KL46 | in database | VP160552 | this study | GCA_905331715 | 1 | 1 | n.a. | | |
| 47 | KL47 | in database | 20542 | this study | GCA_905331615 | 2 | 2 | 2 | | |
| 48 | KL48 | in database | VP120269 | this study | GCA_905331665 | 2 | 2 | 0 | | |
| 49 | KL49 | in database | GCA_002154055 | NCBI | | 2 | 1 | 1 | | |
| 50 | KL50 | in database | 20545 | this study | GCA_905331595 | 1 | 1 | n.a. | | |
| 51 | KL51 | in database | VP120841 | this study | GCA_905331695 | 1 | 1 | n.a. | | |
| 52 | KL52 | in database | VP180273 | this study | GCA_905331905 | 1 | 1 | n.a. | | |
| 53 | KL53 | in database | GCA_001728135 | NCBI | | 2 | 1 | 0 | | |
| 54 | KL54 | in database | GCA_000958585 | NCBI | | 1 | 1 | n.a. | | |
| 55 | KL55 | in database | VP180179 | this study | GCA_905331915 | 1 | 1 | 1 | | |
| 56 | KL56 | in database | VP161624 | this study | GCA_905331825 | 14 | 13 | 12 | | |
| 57 | KL57 | in database | GCA_002150155 | NCBI | | 3 | 3 | n.a. | | |
| 58 | KL58 | in database | GCA_001609345 | NCBI | | 5 | 5 | 2 | | |
| 59 | KL59 | in database | VP162119 | this study | GCA_905331845 | 1 | 1 | n.a. | | |
| 60 | KL60 | in database | VP161546 | this study | GCA_905331765 | 3 | 2 | 1 | | |
| 61 | KL61 | in database | 20556 | this study | GCA_905331605 | 1 | 1 | n.a. | | |
| 62 | | does not exist | | | | | | | | |
| 63 | KL63 | in database | VP162129 | this study | GCA_905331855 | 9 | 8 | 6 | | |
| 64 | KL64 | in database | GCA_001856035 | NCBI | | 2 | 1 | n.a. | | |
| 65 | KL65 | in database | 20564 | this study | GCA_905331625 | 1 | 1 | n.a. | | |
| 66 | KL66 | in database | 20565 | this study | GCA_905331645 | 1 | 1 | n.a. | | |
| 67 | KL67 | in database | 20566 | this study | GCA_905331635 | 1 | 1 | n.a. | | |
| 68 | KL68 | in database | GCA_001273575 | NCBI | | 8 | 8 | 4 | | |
| 68 | KL68-1 | in database | GCA_001728625 | NCBI | | | | | | |
| 69 | KL69 | in database | GCA_000491275 | NCBI | | 4 | 3 | 1 | | |
| 70 | KL70 | in database | GCA_001726855 | NCBI | | 2 | 2 | 2 | | |
| 71 | KL71 | in database | VP32 | CNGB | CNA0007063 | 1 | 1 | 1 | | |

**Serotype UnTypable**

| | | | | | |
|---|---|---|---|---|---|
| KUT1 | KLUT1 | in database | GCA_001584415 | NCBI | |
| KUT2 | KLUT2 | in database | GCA_001609575 | NCBI | |
| KUT3 | KLUT3 | in database | VP162105 | this study | GCA_905331815.1 |
| KUT4 | KLUT4 | in database | VP162125 | this study | GCA_905331835.1 |
| | KLUT4-1 | in database | GCA_001726495 | NCBI | |
| KUT5 | KLUT5 | in database | VP162180 | this study | GCA_905331885.1 |
| KUT6 | KLUT6 | in database | VP161168 | this study | GCA_905331775.1 |
| KUT7 | KLUT7 | in database | VP160744 | this study | GCA_905331735.1 |
| KUT8 | KLUT8 | in database | VP160152 | this study | GCA_905331705.1 |

**Serotype Unknown/New**

| | | | | |
|---|---|---|---|---|
| | KL101 | in database | GCA_000489655 | NCBI |
| | KL102 | in database | GCA_002221185 | NCBI |

| | | | |
|---|---|---|---|
| KL103 | in database | GCA_000490115 | NCBI |
| KL104 | in database | GCA_000500545 | NCBI |
| KL105 | in database | GCA_000523375 | NCBI |
| KL106 | in database | GCA_000591555 | NCBI |
| KL107 | in database | GCA_000707085 | NCBI |
| KL108 | in database | GCA_000732995 | NCBI |
| KL109 | in database | GCA_000786835 | NCBI |
| KL110 | in database | GCA_000877625 | NCBI |
| KL111 | in database | GCA_000972025 | NCBI |
| KL112 | in database | GCA_001268005 | NCBI |
| KL113 | in database | GCA_001726435 | NCBI |
| KL114 | in database | GCA_001268015 | NCBI |
| KL115 | in database | GCA_001610595 | NCBI |
| KL116 | in database | GCA_001273635 | NCBI |
| KL117 | in database | GCA_001433415 | NCBI |
| KL118 | in database | GCA_001608855 | NCBI |
| KL119 | in database | GCA_001727295 | NCBI |
| KL120 | in database | GCA_001727345 | NCBI |
| KL121 | in database | GCA_001727045 | NCBI |
| KL122 | in database | GCA_001727155 | NCBI |
| KL123 | in database | GCA_001727645 | NCBI |
| KL124 | in database | GCA_002018725 | NCBI |
| KL125 | in database | GCA_002150335 | NCBI |
| KL126 | in database | GCA_002153895 | NCBI |
| KL127 | in database | GCA_002153975 | NCBI |
| KL128 | in database | GCA_002154015 | NCBI |
| KL129 | in database | GCA_002018535 | NCBI |
| KL130 | in database | GCA_002144585 | NCBI |
| KL131 | in database | GCA_002221065 | NCBI |
| KL132 | in database | GCA_002221165 | NCBI |
| KL133 | in database | 670.938 | PATRIC |
| KL134 | in database | 670.90 | PATRIC |
| KL135 | in database | 670.812 | PATRIC |
| KL136 | in database | 1620393.3 | PATRIC |
| KL137 | in database | 670.467 | PATRIC |
| KL138 | in database | 670.466 | PATRIC |
| KL139 | in database | 670.2224 | PATRIC |
| KL140 | in database | 670.2093 | PATRIC |
| KL141 | in database | 670.2085 | PATRIC |
| KL142 | in database | 670.2080 | PATRIC |
| KL143 | in database | 670.2053 | PATRIC |
| KL144 | in database | 670.2033 | PATRIC |
| KL145 | in database | 670.1966 | PATRIC |
| KL146 | in database | 670.1961 | PATRIC |
| KL147 | in database | 670.1931 | PATRIC |
| KL148 | in database | 670.1914 | PATRIC |
| KL149 | in database | 670.1676 | PATRIC |
| KL150 | in database | 670.1673 | PATRIC |
| KL151 | in database | 670.1672 | PATRIC |
| KL152 | in database | 670.1657 | PATRIC |
| KL153 | in database | 670.1645 | PATRIC |
| KL154 | in database | 670.1383 | PATRIC |
| KL155 | in database | 670.1377 | PATRIC |
| KL156 | in database | 670.1351 | PATRIC |
| KL157 | in database | 670.1111 | PATRIC |
