## Supplemental Table 2 for "Development of *Kaptive* databases for *Vibrio parahaemolyticus* O- and K-antigen serotyping"

| Strain | Accession | K serogroup | Kaptive | Kaptive match confidence | VPSero |
|---|---|---|---|---|---|
| VP199 | CNA0006925 | K1 | KL1 | Very high | K1 |
| VP107 | CNA0006832 | K11 | KL11 | Good | K11 |
| VP179 | CNA0006903 | K12 | KL12 | High | K12 |
| VP179 | CNA0006903 | K12 | KL12 | High | K12 |
| VP109 | CNA0006834 | K13 | KL13 | Good | K13 |
| VP43 | CNA0007181 | K17 | KL17 | Good | K17 |
| VP43 | CNA0007181 | K17 | KL17 | Good | K17 |
| VP197 | CNA0006923 | K18 | KL18 | Good | K18 |
| VP197 | CNA0006923 | K18 | KL18 | Good | K18 |
| VP192 | CNA0006918 | K19 | KL19 | High | K19 |
| VP323 | CNA0007056 | K19 | KL19 | High | Knt |
| VP206 | CNA0006934 | K20 | KL20 | Low | Knt |
| VP207 | CNA0006935 | K20 | KL20 | Good | Knt |
| VP439 | CNA0007180 | K21 | KL21 | Good | K21 |
| VP202 | CNA0006930 | K23 | KL23 | Good | K23 |
| VP202 | CNA0006930 | K23 | KL23 | Good | K23 |
| VP132 | CNA0006860 | K25 | KL25 | Low | K25 |
| VP132 | CNA0006860 | K25 | KL25 | Low | K25 |
| VP53 | CNA0007199 | K28 | KL28 | High | K28 |
| VP247 | CNA0006976 | K29 | KL29 | None | K29 |
| VP245 | CNA0006974 | K3 | KL3 | Good | K3 |
| VP190 | CNA0006916 | K30 | KL30 | Good | K30 |
| VP190 | CNA0006916 | K30 | KL30 | Good | K30 |
| VP205 | CNA0006933 | K31 | KL31 | Good | K31 |
| VP99 | CNA0007243 | K32 | KL32 | Good | K32 |
| VP99 | CNA0007243 | K32 | KL32 | Good | K32 |
| VP203 | CNA0006931 | K33 | KL33 | Good | K33 |
| VP6 | CNA0007217 | K34 | KL34 | Good | K34 |
| VP104 | CNA0006830 | K36 | KL36 | Good | K36 |
| VP122 | CNA0006849 | K36 | KL36 | None | Knt |
| VP196 | CNA0006922 | K37 | KL37 | None | Knt |
| VP239 | CNA0006969 | K37 | KL124 | Good | K37 |
| VP200 | CNA0006928 | K38 | KL38 | Good | K38 |
| VP198 | CNA0006924 | K4 | KL4 | Good | K4 |
| VP234 | CNA0006964 | K41 | KL41 | Good | K41 |
| VP4 | CNA0007195 | K42 | KL42 | Good | K42 |
| VP4 | CNA0007195 | K42 | KL42 | Good | K42 |
| VP113 | CNA0006839 | K44 | KL44 | Good | K44 |
| VP229 | CNA0006958 | K48 | KL48 | None | K48 |
| VP238 | CNA0006968 | K49 | KL67 | None | K49 |
| VP195 | CNA0006921 | K5 | KL5 | Good | K5 |
| VP1 | CNA0006927 | K55 | KL55 | Good | K55 |
| VP1 | CNA0006927 | K55 | KL55 | Good | K55 |
| VP334 | CNA0007068 | K56 | KL56 | Good | K56 |
| VP334 | CNA0007068 | K56 | KL56 | Good | K56 |
| VP230 | CNA0006960 | K58 | KL6 | High | K6 |

| | | | | | |
|---|---|---|---|---|---|
| VP204 | CNA0006932 | K6 | KL6 | High | K6 |
| VP204 | CNA0006932 | K6 | KL6 | High | K6 |
| VP16 | CNA0006893 | K60 | KL60 | Perfect | K60 |
| VP321 | CNA0007054 | K63 | KL63 | Low | K63 |
| VP321 | CNA0007054 | K63 | KL63 | Low | K63 |
| VP161 | CNA0006887 | K68 | KL68-1 | None | K68 |
| VP33 | CNA0007074 | K69 | KL69 | Very high | K69 |
| VP32 | CNA0007063 | K71 | KL71 | Good | K71 |
| VP187 | CNA0006912 | K8 | KL8 | Good | K8 |
| VP135 | CNA0006863 | K9 | KL9 | Good | K9 |
| VP135 | CNA0006863 | K9 | KL9 | Good | K9 |
| VP127 | CNA0006854 | KUT | KL66 | Good | Knt |
| VP12 | CNA0006857 | KUT | KL32 | Good | K32 |
| VP13 | CNA0006868 | KUT | KL102 | Good | Knt |
| VP153 | CNA0006882 | KUT | KL24 | Very high | Knt |
| VP189 | CNA0006914 | KUT | KL54 | Low | Knt |
| VP18 | CNA0006915 | KUT | KL42 | Good | K42 |
| VP22 | CNA0006959 | KUT | KL6 | High | K6 |
| VP236 | CNA0006966 | KUT | KL140 | High | Knt |
| VP240 | CNA0006971 | KUT | KL22 | Good | Knt |
| VP241 | CNA0006972 | KUT | KL50 | High | Knt |
| VP257 | CNA0006987 | KUT | KL67 | None | Knt |
| VP262 | CNA0006992 | KUT | KL67 | None | Knt |
| VP264 | CNA0006994 | KUT | KL6 | High | K6 |
| VP269 | CNA0006999 | KUT | KL22 | Good | Knt |
| VP271 | CNA0007000 | KUT | KL22 | Good | Knt |
| VP285 | CNA0007015 | KUT | KL6 | High | K6 |
| VP287 | CNA0007017 | KUT | KL101 | None | Knt |
| VP300 | CNA0007033 | KUT | KL110 | Low | Knt |
| VP301 | CNA0007034 | KUT | KL24 | Very high | Knt |
| VP303 | CNA0007036 | KUT | KL70 | Good | K70 |
| VP306 | CNA0007039 | KUT | KL7 | Good | Knt |
| VP308 | CNA0007041 | KUT | KL52 | None | Knt |
| VP313 | CNA0007046 | KUT | KL145 | Good | Knt |
| VP315 | CNA0007048 | KUT | KL43 | None | Knt |
| VP318 | CNA0007050 | KUT | KL7 | Good | Knt |
| VP344 | CNA0007079 | KUT | KLUT5 | Good | Knt |
| VP351 | CNA0007086 | KUT | KL126 | Good | Knt |
| VP352 | CNA0007087 | KUT | KL134 | None | Knt |
| VP355 | CNA0007090 | KUT | KL119 | Low | Knt |
| VP356 | CNA0007091 | KUT | KL156 | High | Knt |
| VP358 | CNA0007093 | KUT | KL147 | Low | Knt |
| VP360 | CNA0007096 | KUT | KL126 | None | Knt |
| VP361 | CNA0007097 | KUT | KL123 | Good | Knt |
| VP362 | CNA0007098 | KUT | KL123 | Good | Knt |
| VP364 | CNA0007100 | KUT | KL22 | Good | Knt |
| VP365 | CNA0007101 | KUT | KL6 | High | K6 |

| | | | | | |
|---|---|---|---|---|---|
| VP366 | CNA0007102 | KUT | KL114 | None | Knt |
| VP368 | CNA0007103 | KUT | KL45 | Low | Knt |
| VP371 | CNA0007106 | KUT | KL43 | None | Knt |
| VP374 | CNA0007109 | KUT | KL6 | High | K6 |
| VP379 | CNA0007113 | KUT | KLUT5 | Good | Knt |
| VP382 | CNA0007117 | KUT | KL149 | Good | Knt |
| VP383 | CNA0007118 | KUT | KLUT5 | Good | Knt |
| VP385 | CNA0007120 | KUT | KLUT3 | Good | Knt |
| VP386 | CNA0007121 | KUT | KL111 | Good | Knt |
| VP387 | CNA0007122 | KUT | KL124 | None | Knt |
| VP388 | CNA0007123 | KUT | KL112 | None | Knt |
| VP389 | CNA0007124 | KUT | KL107 | Good | Knt |
| VP393 | CNA0007129 | KUT | KL32 | Good | K32 |
| VP395 | CNA0007131 | KUT | KL6 | High | K6 |
| VP39 | CNA0007136 | KUT | KL110 | Low | Knt |
| VP400 | CNA0007138 | KUT | KL123 | Good | Knt |
| VP403 | CNA0007141 | KUT | KL6 | High | K6 |
| VP408 | CNA0007146 | KUT | KLUT3 | Good | Knt |
| VP409 | CNA0007147 | KUT | KLUT3 | Good | Knt |
| VP410 | CNA0007149 | KUT | KLUT5 | Good | Knt |
| VP411 | CNA0007150 | KUT | KLUT3 | Good | Knt |
| VP412 | CNA0007151 | KUT | KLUT3 | Good | Knt |
| VP413 | CNA0007152 | KUT | KLUT3 | Good | Knt |
| VP414 | CNA0007153 | KUT | KLUT3 | Good | Knt |
| VP415 | CNA0007154 | KUT | KLUT3 | Good | Knt |
| VP417 | CNA0007156 | KUT | KLUT5 | Good | Knt |
| VP418 | CNA0007157 | KUT | KLUT5 | Good | Knt |
| VP419 | CNA0007158 | KUT | KLUT3 | Good | Knt |
| VP420 | CNA0007160 | KUT | KLUT5 | Good | Knt |
| VP422 | CNA0007162 | KUT | KLUT5 | Good | Knt |
| VP423 | CNA0007163 | KUT | KLUT5 | Good | Knt |
| VP424 | CNA0007164 | KUT | KLUT5 | Good | Knt |
| VP425 | CNA0007165 | KUT | KLUT3 | Good | Knt |
| VP426 | CNA0007166 | KUT | KLUT3 | Good | Knt |
| VP427 | CNA0007167 | KUT | KLUT3 | Good | Knt |
| VP429 | CNA0007169 | KUT | KL118 | Good | Knt |
| VP430 | CNA0007171 | KUT | KLUT5 | Good | Knt |
| VP431 | CNA0007172 | KUT | KLUT5 | Good | Knt |
| VP432 | CNA0007173 | KUT | KLUT5 | Good | Knt |
| VP433 | CNA0007174 | KUT | KL64 | None | Knt |
| VP435 | CNA0007176 | KUT | KLUT5 | Good | Knt |
| VP436 | CNA0007177 | KUT | KLUT3 | Good | Knt |
| VP437 | CNA0007178 | KUT | KLUT5 | Good | Knt |
| VP438 | CNA0007179 | KUT | KLUT5 | Good | Knt |
| VP445 | CNA0007187 | KUT | KLUT3 | Good | Knt |
| VP446 | CNA0007188 | KUT | KLUT2 | Low | Knt |
| VP46 | CNA0007191 | KUT | KL22 | Good | Knt |

| | | | | | |
|---|---|---|---|---|---|
| VP47 | CNA0007192 | KUT | KL22 | Good | Knt |
| VP52 | CNA0007198 | KUT | KL37 | None | Knt |
| VP5 | CNA0007206 | KUT | KL25 | None | Knt |
| VP62 | CNA0007209 | KUT | KL46 | Good | Knt |
| VP68 | CNA0007215 | KUT | KL43 | None | Knt |
